# Collagen binding adhesin restricts *Staphylococcus aureus* skin infection

**DOI:** 10.1101/2024.11.01.621145

**Authors:** Mohini Bhattacharya, Brady L. Spencer, Jakub M. Kwiecinski, Magdalena Podkowik, Gregory Putzel, Alejandro Pironti, Bo Shopsin, Kelly S. Doran, Alexander R. Horswill

## Abstract

*Staphylococcus aureus* causes approximately 80% of skin and soft tissue infections (SSTIs). Collagen is the most abundant human extracellular matrix protein with critical roles in wound healing, and *S. aureus* encodes a collagen binding adhesin (Cna). The role of this protein during skin infections is unknown. Here we report that inability to bind collagen results in worsened pathology of intradermal Δ*cna S. aureus* infection. WT/Cna+ *S. aureus* showed reduced infection severity, aggregate formation, and significantly improved clearance of bacteria. Cna binds to the collagen-like domain of serum C1q protein to reduce its opsonophagocytic functions. We demonstrate that infection of C1qKO mice with WT bacteria show results similar to the Δ*cna* group. Conversely, inability to bind collagen resulted in an amplified inflammatory response caused in part by macrophage and neutrophil small molecule mediators released at the infection site (MMP-9, MMP-12, LTB_4_), resulting in increased immune cell infiltration and death.

## Introduction

Collagen is the most abundant component of the human extracellular matrix, forming 30% of the protein dry weight in the human body^1^. With an essential role in wound healing, collagen is made up of 3 polypeptide alpha chains with characteristic Gly-X-X’ motifs that assemble to form a right-handed helical structure commonly identified as the ‘collagen like domain’^2,3^. *Staphylococcus aureus* is the most common cause of <u>s</u>kin and <u>s</u>oft tissue infections (SSTIs) such as abscesses, carbuncles and furuncles^4,5^. The treatment of these infections is often complicated by the acquisition of antibiotic resistance, the most common of which is the development of <u>m</u>ethicillin resistant *<u>S</u>. <u>a</u>ureus* (MRSA)^5,6^. USA300 is a particularly problematic MRSA that has been implicated in a large percentage of SSTIs in the Unites States, for more than a decade^6,7^. The success of *S. aureus* as a pathogen is in part because it is particularly adept at expressing numerous toxins that target host cells or evade immune responses^8,9^. Once such evasion tactic is the expression of surface adhesins that bind host extracellular matrix components^10^. Among these proteins is the <u>c</u>ollagen bi<u>n</u>ding <u>a</u>dhesin (Cna) expressed by a subset of strains. USA400 strain MW2, is another epidemic MRSA with steadily reducing incidences in the United States, Europe and Canada over the past 15 years^11–14^. MW2 expresses Cna. As a typical sortase anchored protein, Cna is attached to the cell wall via a C-terminal LPXTG motif^15^. The N-terminal A domain is characterized as being required for binding to collagen in the *S. aureus* strain Phillips^16^. Multiple mouse models of intravenous injection demonstrate a role for Cna in the virulence of septic arthritis, keratitis and osteomyelitis caused by *S. aureus,* some with conflicting results^17–21^. There are currently no documented roles for Cna in the pathogenesis of *S. aureus* skin infection.

The N-terminal A domain of Cna is also reported to bind the N-terminal collagen-like tail of serum protein C1q to prevent the classical pathway of complement mediated killing in strain Phillips^22^. While the C-terminal globular head of C1q binds either directly to bacteria via pathogen associated molecular patterns (PAMPs), or to immunoglobulins on the bacterial surface, the N-terminal collagen like domain is recognized by innate immune cells^23^. Neutrophils and macrophages are key effectors in the phagocytic removal of opsonized bacteria. Following recognition of C1q bound to bacteria, a series of host proteolytic events results in the formation of C3b, via the C3 convertase (C4b2a). Deposition of C3b on the surface results in bacterial uptake by neutrophils and macrophages which utilize toxic, antibacterial effectors to eliminate bacterial populations from the infection site^24^. Alternatively, immune cells that are unable to phagocytose bacteria will use degranulation, reactive oxygen burst and extracellular trap formation as mechanisms to release antimicrobial compounds that kill bacteria^25,26^. Unlike phagocytosis, these processes can result in a higher degree of inflammation. Matrix metalloproteases (MMPs) are an important subclass of soluble mediators, with Zn^2+^ and Ca^2+^ dependent endopeptidase activities required for tissue remodeling. MMP-9 is released largely by neutrophils during degranulation but also by monocytes and macrophages. In addition to its collagenase activity, MMP-9 enhances TNF-α and IL-1β signaling and binds to CXCL8, cleaving the cytokine to increase its chemotactic potency 10-fold^27^. Additionally, the collagenase activity of MMP-9 results in the formation of the inflammatory fragment, Pro-Gly-Pro which is also a potent neutrophil chemotactic factor, thereby augmenting the inflammatory response to infection. Multiple reports demonstrate that MMP-9 is a significant contributing factor for pathogen control and removal^28–30^. The elastase MMP-12 is released primarily from macrophages and is required for macrophage infiltration into infection sites. The hemopexin domain of this protein is reported to have bactericidal activity specifically against *S. aureus* ^31,32^. The coordinated activity of neutrophil and macrophage derived MMPs cause the activation of leukotriene hydrolases such as LTAH_4_ which results in the release of leukotriene B_4_, an additional pro-inflammatory mediator^33^. The contribution of these MMPs at the infection site is therefore significant to the inflammatory cascade induced by infection.

Our studies demonstrate that Cna+ *S. aureus* causes skin infections with significantly reduced bacterial loads accompanied by less inflammation, when compared with those caused by *S. aureus* unable to express Cna. We show that binding to collagen and the collagen-like motif of C1q reduces bacterial spread and the inflammatory response to infection respectively. The two major populations of immune cells affected by Cna-expressing bacteria are neutrophils and macrophages, both of which instigate a cycle of inflammation at the infection site that is propagated by increased release of matrix metalloproteases. Altogether this is the first report for the role of Cna in *S. aureus* skin infection and highlights the significance of collagen to infection outcomes.

## Results

### Expression of collagen binding adhesin (Cna) is sufficient to limit *S. aureus* abscess formation and bacterial burden

To evaluate the role of collagen binding adhesin in the outcome of *S. aureus*-associated skin infections, we utilized the clinical, MRSA USA400 MW2 strain of *S. aureus* that expresses Cna (hereafter WT), an isogenic mutant unable to synthesize Cna*, (*Δ*cna*) and a corresponding Δ*cna* strain where *cna* is ectopically expressed from a plasmid Δ*cna:pCna* (hereafter comp *cna*). *In vitro* adhesion assays demonstrate that expression of Cna by WT MW2 is sufficient for bacterial binding to collagen, a phenotype that is abrogated in the Δ*cna* isogenic mutant (**Figure S1A**). Collagen binding was restored in comp *cna* bacteria and *cna* was required to induce binding when heterologously expressed in the non-pathogenic *Staphylococcus carnosus* (**Figure S1B**).

We therefore utilized this MRSA strain in an established model of intradermal abscess infection and monitored abscess size and weight loss in female, age matched BALB/c mice over a 7-day period^34,35^. At the end of the experiment, lesions were excised and <u>c</u>olony forming <u>u</u>nits per <u>g</u>ram (CFU/g) of homogenized tissue were measured (**Figure 1A**). WT and comp *cna* infected animals demonstrated weight loss comparable to the negative control group injected with saline, while Δ*cna* infected mice lost ∼10-20% of their initial weight (**Figure 1B**). Mice infected with the Δ*cna* mutant also formed discernably larger abscesses when compared with WT and comp *cna* infected groups (**Figure 1C, D**). Larger lesions were accompanied by higher levels of recovered CFU/g (∼1.5 log) of tissue in these mice, indicating exacerbated skin abscess infection in the absence of Cna (**Figure 1E**).

**Figure 1.**
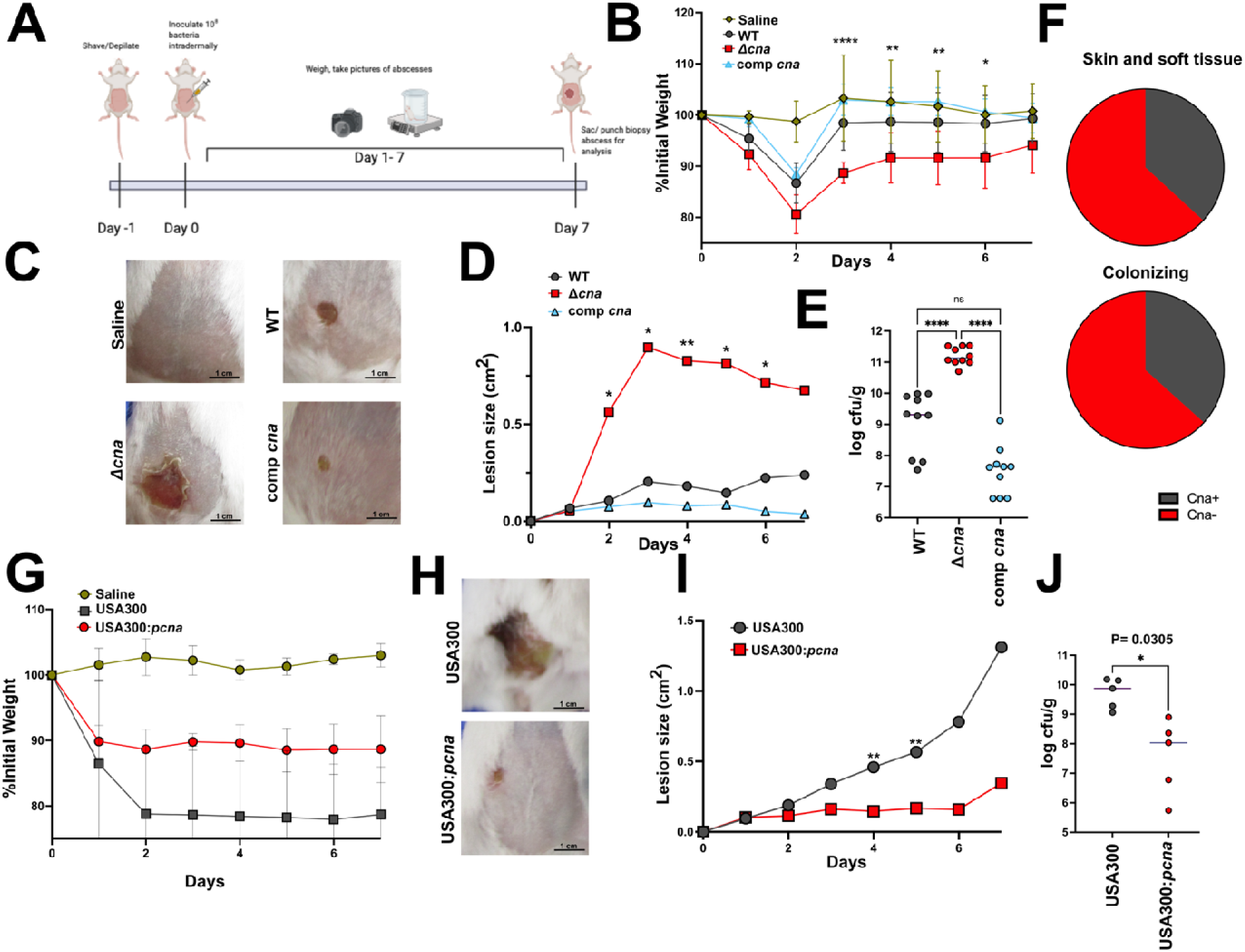
Collagen binding adhesin reduces severity of *S. aureus* skin infection. Method used for intradermal *S. aureus* infection and abscess formation (**A**). Weight loss measured over the 7-day infection period in mice (n=10 per bacterial strain) infected with WT MW2, isogenic Δ*cna* or comp *cna* bacteria and calculated as a percentage of values at day 1(**B**). Images of abscess lesions taken at day 7, representative of mice infected with strains mentioned in B (**C**). Measurement of lesion sizes from mice infected with strains as above over a period of 7 days. Measurement were made using Image J (**D**). Bacterial burdens enumerated per gram of homogenized tissue excised at day 7 post infection with strains as described above (**E**). Percent distribution of the *cna* gene as detected in *S. aureus* clones associated with human skin and soft tissue infections (n=20) and colonizing healthy anterior nares (n=30) (**F**) Abscess model of skin infection performed as described in A, to compare pathology caused by USA300 (Cna-) an USA300:*pcna* (Cna+) bacteria (n=5 per group). Weight loss was measured over 7 days and calculated as percentage of values at day 0 (**G**). Representative images of lesions formed 7 days post infection with strains mentioned in G (**H**). Lesion sizes measured over 7 days post infection with strains as described in G. Measurements were made using Image J (**I**). Colony forming units (CFU) per gram of homogenized tissue enumerated from abscesses biopsied from mice infected with strains as mentioned in G, at day 7 post inoculation (**J**). Results are representative of 3 (MW2) and 2 (USA300) independent analyses. Statistical analyses were performed with a one-way ANOVA (E) a two tailed Students t-test (I) or a two-way ANOVA with Bonferroni post test. Error was calculate based on SEM. Groups with significant differences are denoted. *P<0.05, **P<0.01, ****P<0.0001.

To gauge the impact of Cna to clinical infections, we assessed the prevalence of the *cna* gene in *S. aureus* isolates representing the major clonal types from patients with primary SSTIs^36^. Of note, 60% of these isolates did not encode for *cna.* Since nasal colonization is an important risk factor for infection, we also sequenced for the presence of the *cna* gene across the major clonal types in patients with positive nasal cultures for *S. aureus*^37^. We similarly found that ∼60% of these isolates did not encode for *cna* (**Figure 1F**). These results indicate that the absence of Cna is common and, depending on the population, may predominate in both nasal colonization and SSTIs associated with *S. aureus*.

We performed a mouse intradermal infection using USA300, a dominant clinical strain of MRSA which does not encode the *cna* gene^38^. We infected mice as mentioned above and compared the progression of abscess formation with a USA300 strain engineered to express Cna from the vector used above (USA300:*pCna*). As expected, we found that USA300 (Cna-) infected mice showed ∼10% higher rates of weight loss when compared to the USA300:*pCna* group (**Figure 1G**). This was comparable to observations made in mice infected with the Δ*cna* isogenic mutant in the MW2 WT background. Similarly, we observed macroscopically larger abscesses, which was confirmed by measuring lesion sizes over the course of infection (**Figure 1H**, **I**). Lastly, these phenotypes were associated with ∼2 log higher CFU/g of tissue in USA300 infected animals when compared to the USA300:*pCna* group (**Figure 1J**).

Finally, to verify that these phenotypes were not due to sex differences in our mouse experiments, we performed a similar infection study in male, age matched BALB/c mice. Consistent with the results from experiments with female mice of the same background, we found that male BALB/c mice infected with Δ*cna* bacteria had visibly larger abscesses, with larger lesions sizes measured through the course of infection (**Figure S1C**). While we did not observe significant weight loss, bacterial burdens were similarly ∼1-1.5 log higher in Δ*cna* infected mice, as compared with WT and comp *cna* infected controls (**Figure S1D, E**). Collectively these data confirm that the expression of Cna is sufficient to restrict the bacterial burden and gross pathology observed during *S. aureus* skin infection.

### Collagen binding dampens the inflammatory response to *S. aureus* in skin abscesses

*S. aureus* skin infections are often associated with an inflammatory immune response that causes worsened pathology^39^. <u>H</u>ematoxylin <u>E</u>osin (H&E) staining performed on longitudinal sections of abscess tissue excised at day 7 determined that the worsening of infection phenotypes we observed were indeed associated with the marked accumulation of necrotic tissue (purple) in Δ*cna* infected animals when compared to those infected with either WT or comp *cna S. aureus,* which were more comparable to control animals injected with saline (pink) (**Figure 2A**). We therefore utilized a multiplex assay to measure the concentrations of 44 inflammatory cytokines in these tissue samples. We observed a comprehensive increase in the inflammatory response measured from Δ*cna* infected mice (**Figure 2B**). Specifically, this included significant increases in the concentrations of key markers of inflammation namely IL-6, TNF-a and IL-1b as well as chemotactic factors for hemopoietic cells, KC, MCP-1, G-CSF and GM-CSF. Additionally, infection with Δ*cna* bacteria caused a significant increase in concentration of the tissue inhibitor of <u>m</u>atrix metallo<u>p</u>rotease-1 (TIMP-1) (**Figure 2C**). Collectively these results indicate that Δ*cna* bacteria induce a larger skin inflammatory response when compared with an isogenic Cna+ counterpart.

**Figure 2.**
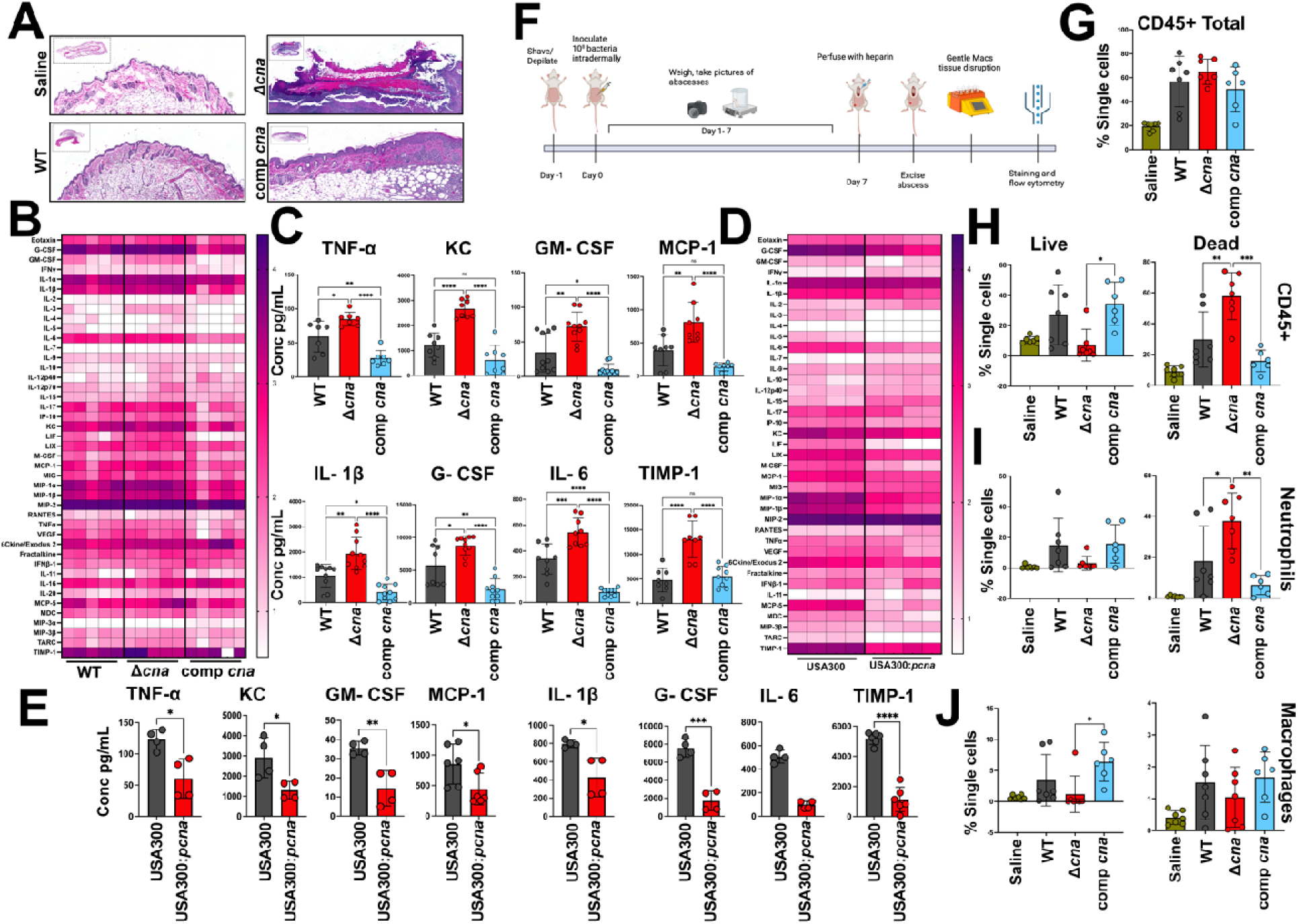
Expression of collagen binding adhesin is sufficient to restrict host inflammatory response. Hematoxylin Eosin staining of tissue sections biopsied from mice infected with WT, isogenic Δ*cna* or comp *cna* bacteria at day 7 post infection. Tissue section of mouse injected with saline is shown as a negative control (**A**). Cytokine array performed on mouse tissue collected from animals infected as described for A. The multiplexin analysis was performed to measure the concentration of 44 cytokines, using the Luminex™ 200 system by Eve Technologies Corp. Each column represents results from a single mouse (n=5 per group) (**B**). Individual graphs t show differences in cytokine measurements as made in B, for 8 cytokines of interest generated from mice i response to respective bacterial strains (**C**). Quantification of cytokine concentrations similar to B, made from mic infected with USA300 or the isogenic USA300:*pcna* strain (n=5 per group) (**D**). Individual graphs shown for cytokines similar to C, measured from abscesses infected with strains as mentioned in D, 7 days post bacterial inoculation (**E**). Summary of methods used to perform flow cytometry quantification of immune cells from abscesses infected with bacteria as described for A (**F**). Quantification of the total number of single immune cells using a antibody specific to CD45, from tissue samples collected as described in F, for mice infected with bacterial strains described in A (n=7) (**G**). Differentiation of cells enumerated in G, based on exclusion of Am Cyan viability dye(live) from observably dead populations (**H**). Sub populations of total CD45 cells classified as neutrophils (**I**) or macrophages (**J**) based on staining with cell specific antibodies as described in Figure S2A, B. Results ar representative of 3 independent analyses. Statistical analyses were performed with a one-way ANOVA with a Bonferroni post-test. Error was calculated based on SEM. *P<0.05, **P<0.01, ****P<0.0001

Since *S. aureus* expresses multiple toxins that can cause host immune cell lysis and acute inflammation, we wanted to confirm that Cna was sufficient to induce these changes in the host. We therefore performed a similar cytokine analysis using the dominant clinical strain, USA300 which does not express Cna. These measurements were performed in comparison to a USA300:*pCna*, as mentioned for **Figure 1**. USA300, much like the MW2 Δ*cna* infected mice, showed an overall increase in levels of inflammatory cytokines compared to mice infected with USA300:*pCna* bacteria (**Figure 2D, E**).

Alpha toxin is a well-documented dermonecrotic protein expressed by most strains of *S. aureus* ^40,41^. To confirm that the phenotypes we observed in mice are not due to the expression of alpha toxin, we performed an intradermal infection as described above, and compared a transposon mutant of *hla,* the gene encoding the alpha hemolysin protein, with an isogenic *hla::Tn* Δ*cna* strain in the MW2 strain background. Similar to previous results, the inability to express Cna caused an increase in lesion size, weight loss and CFU burden in the *hla::Tn* background, indicating that the phenotypes observed were largely independent of alpha toxin (**Figure S1F-H**). Collectively, these results demonstrate that the expression of Cna is sufficient to suppress the acute intradermal inflammatory response to *S. aureus* infection.

### Absence of Cna results in excessive immune cell death

To quantify immune populations that were contributing to increased inflammation in mice infected with Δ*cna* bacteria, we performed flow cytometry on perfused abscess tissue collected at day 7, using previously published methods^42^ (**Figure 2F, Figure S2A, B**). While the total numbers of CD45^+^ immune cells (as a proportion of total single cells) were comparable between WT, Δ*cna* and comp *cna* infected groups of mice (**Figure 2G**), we observed a large disparity in the numbers of live, and dead/dying CD45^+^ populations between these groups, with Δ*cna* infected tissue samples containing very few live CD45^+^ cells and a large population of observable dead CD45^+^ cells compared to WT infected samples (**Figure 2H**). This phenotype was able to be complemented, as numbers of both live and dead CD45+ cells could be restored to WT levels in mice infected with the comp *cna* strain. These data indicate that the loss of Cna results in abscesses composed primarily of dead immune cells. Further analysis demonstrated that the observed increase in dead CD45^+^ cells in Δ*cna* infected animals was driven largely by an increase in (intact but) dead neutrophils (**Figure 2I**). Interestingly, we did not observe this increase of dead cells within the macrophage population **(Figure 2J)**. Inversely, comparison of Δ*cna* to WT and especially comp *cna* infected groups confirmed that expression of Cna may promote survival of neutrophils (**Figure 2I**) and macrophages (**Figure 2J**), since live populations of these cells were restored to significantly higher levels in comp *cna* infected tissue. Although to a smaller extent, this decrease in live immune cells was also observed for additional CD45+ populations (**Figure S2C**). Collectively these results demonstrate a rise in total immune cell death, particularly neutrophils, during infection with *S. aureus* lacking Cna.

In an attempt to avoid the extensive cell death observed at day 7, we assessed the immune profile of these infections at an earlier time point. Our results indicate that Δ*cna* infected mice develop larger abscesses as early as day 3, when compared to WT and comp *cna* groups (**Figure 1D**). H&E staining revealed pathology in Δ*cna* infected mice at day 3 that was similar to that observed at day 7, although the levels of inflammation were less pronounced (**Figure S2D-G**). We observed that gross infection phenotypes were similar to day 7 including weight loss (**Figure S3A**), lesion sizes (**Figure S3B**),CFU burdens (**Figure S3C**) and inflammatory cytokine responses (**Figure S3D, E**).

We next used the previously described antibody panel to quantify immune cell populations present in day-3 abscess tissue, collected and performed as previously described. Similar to results from day 7, while the CD45+ cells (as a proportion of total single cells) were similar among *S. aureus* infected groups, we again observed very few live immune cells in Δ*cna* infected lesion tissue, compared to both WT and comp *cna* infected animals (**Figure S3F**). Concurrently, we observed significantly higher populations of dead/dying cells in Δ*cna* infected abscesses compared to WT and comp *cna* infected abscesses, again largely driven by increases in dead neutrophils (**Figure S3G**). Significantly higher numbers of live macrophages could be observed in WT infected tissues, compared to Δ*cna* samples (**Figure S3H**). Similar to results at day 7, the decrease in total numbers of live cells was also evident in additional subpopulations of CD45^+^ cells (**Figure S3I**). Collectively these results confirm that the absence of Cna results in increased inflammation, likely due to neutrophil death over the course of infection.

### Increased uptake of Δ*cna* bacteria causes neutrophil lysis

Serum complement protein C1q plays an important role in the opsonophagocytosis of *S. aureus*^43^. C1q consists of a C-terminal globular head domain that binds to immunoglobulins or directly to the bacterial surface to activate the classical proteolytic pathway that leads to opsonization and phagocytosis of *S. aureus*^44^. The N-terminus of C1q is a collagenous tail domain that has previously been shown to bind to Cna derived from *S. aureus* strain Phillips *in vitro* and inhibit downstream activation of the classical pathway ^22^(**Figure S4A**). To confirm similar *in vitro* binding of Cna to the C1q N-terminal tail in the MW2 strain background, we utilized a competitive enzyme linked immunosorbent assay with a C1q-coated surface to demonstrate a reciprocal relationship wherein the level of Cna bound to C1q progressively decreased when collagen was incubated in the presence of increasing concentrations of recombinant Cna (**Figure S4B**). Since we observed increases in populations of dead neutrophils during *in vivo* infection **(Figure 2I)**, we sought to examine the effect of this interaction on downstream activation of the classical complement pathway. We opsonized WT, Δ*cna* and comp *cna* and measured bound C3b or C4b. C4b is part of the C3 convertase and is cleaved to activate C3b, an opsonin with a central role in the complement pathway^23,44^. Using flow cytometry, we confirmed that Δ*cna* bacteria exhibited significantly higher levels of C4b (**Figure 3A**) and C3b (**Figure 3B**) deposition, compared with both WT and comp *cna* strains. Consistent with the previous study, these results indicate that the presence of Cna reduces opsonization of bacteria by the classical complement pathway^22^. To assess the effect of this Cna mediated inhibition of C1q specifically on phagocytic activity, we measured bacterial uptake in the presence of primary human neutrophils using previously published methods^45–47^. We observed a significant increase in the uptake of Δ*cna* bacteria (∼20%) when compared to neutrophils exposed to WT or comp *cna* for 10 minutes (**Figure 3C**). To attribute a direct role to C1q in this process, we opsonized bacteria with C1q-depleted serum prior to performing the experiment. We found no significant differences in uptake between WT, Δ*cna* and comp *cna* under these conditions (**Figure 3D**). Collectively these results suggest that Cna binds to serum C1q, reducing complement activation and subsequent phagocytic uptake by neutrophils.

**Figure 3.**
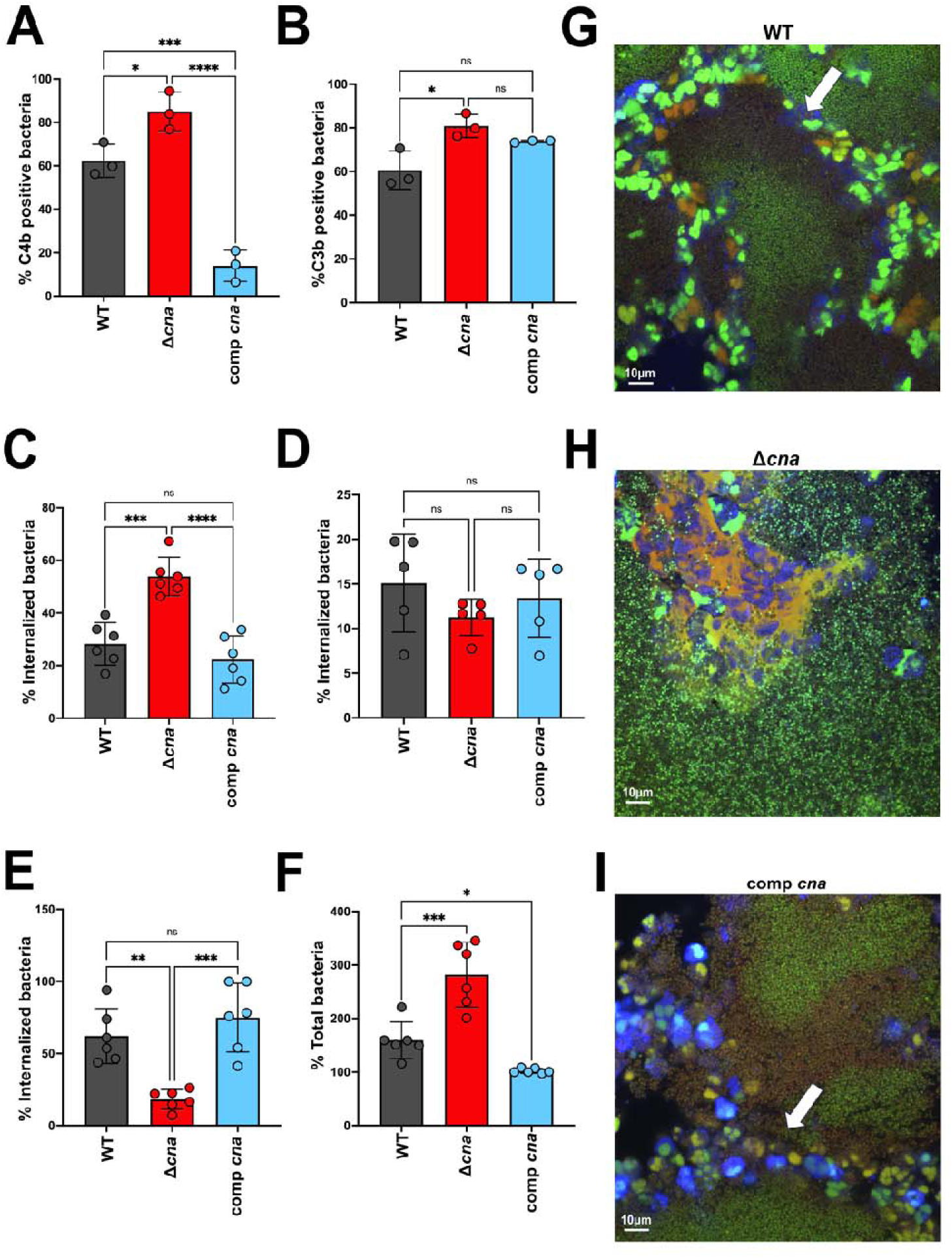
Cna binds to collagen motifs to alter the neutrophil response to bacteria. Flow cytometry of WT, Δ*cn* or comp *cna* bacteria opsonized with 10% serum and stained with antibodies targeting serum complement proteins C4b (**A**) and C3b (**B**). Uptake of bacterial strains mentioned above by primary human neutrophils, following opsonization with 10% pooled human serum. CFU was enumerated 10 minutes post incubation, following which samples were treated with lysostaphin to exclude extracellular populations (**C**). Experiment similar to C performed with bacteria opsonized with 10% C1q-depleted, pooled human serum (**D**). Experiments similar to C, with intracellular (**E**) and total (**F**) bacterial survival calculated at 30 minutes post incubation. Confocal microscopy performed on respective bacterial strains (Green=Syto-9/live) opsonized as mentioned above in the presence of type 1 collage and incubated with Cell Tracker Blue-labelled primary human neutrophils for 20 minutes, following which samples were stained with ethidium homodimer-1 (red) to visualize dead/dying cells. Arrows indicate likely location of collage boundary (**G-I**). Statistical analyses were performed with a one-way ANOVA and Bonferroni post-test. Error was calculated based on SEM. *P<0.05, **P<0.01, ****P<0.0001

We hypothesized that the increase in opsonophagocytic activity observed in response to Δ*cna* results in neutrophil death. We therefore measured bacterial survival 30 minutes post incubation with neutrophils and observed significantly lower numbers of intracellular Δ*cna* bacteria, compared to neutrophils that were exposed to WT or comp *cna S. aureus* (**Figure 3E**). To understand if this was due to neutrophil lysis, bacterial release, and inadvertent killing due to lysostaphin treatment, we measured bacterial survival in the absence of lysostaphin. We observed a significantly larger population of surviving Δ*cna* bacteria compared with both the WT and comp *cna* under these conditions (**Figure 3F**). These observations confirm that the Cna-C1q interaction decreases phagocytic uptake by neutrophils, inadvertently allowing for controlled, efficient killing of the pathogen by these cells. Conversely, in the absence of Cna, we find that neutrophils increasingly phagocytose bacteria which results in cell death and release of Cna-negative *S. aureus*.

### Cna interacts with collagen to restrict neutrophil access to bacteria

Since both collagen and C1q would be present and capable of binding to WT Cna+ bacteria *in vivo*, we performed confocal microscopy on bacteria opsonized in the presence of collagen and incubated with primary human neutrophils as described above. Binding to collagen restricted the direct access of neutrophils to WT (**Figure 3G**) and comp *cna* bacteria (**Figure 3I**). Loss of Syto-9 staining in bacteria closest to the periphery of the collagen-bound aggregate indicated bacterial killing (**Figure 3G**, **I** white arrows). In sharp contrast, Δ*cna* bacteria largely caused lysis of neutrophils as evidenced by ethidium homodimer-1 staining. Similar to results from the opsonophagocytosis assay, we observed large numbers of Syto-9 labeled, extracellular Δ*cna* bacteria, indicating bacterial survival (**Figure 3H**). Together these results indicate that binding to collagen allows for controlled, efficient clearance of bacteria. Conversely, neutrophils have direct access to Δ*cna* bacteria even in the presence of collagen, inadvertently causing them to lyse.

To assess the consequence of the collagen-Cna interaction *in vivo*, we performed multispectral imaging on tissue sections collected 3 days post infection (**Figure 4A**). WT/Cna+ *S. aureus* was observed as DAPI-stained aggregates surrounded by collagen (**Figure 4B**). These were confirmed to be bacteria using a modified Gram stain (**Figure S4C**). We found that macrophages and neutrophils were present but restricted from the infection in WT infected abscesses (**Figure 4B**). In contrast, Δ*cna* infections showed a higher number of neutrophils and decrease in observable macrophages, both in close juxtaposition to bacteria, which were more diffused compared to both WT and comp *cna* infected tissue (**Figure 4C-D**). Quantification of these immune cells in the tissue section corroborated our previous flow cytometry results to show comparable levels of CD45+ populations with higher numbers of neutrophils (∼30%) in Δ*cna* infected samples compared to both WT and comp *cna* sections (**Figure 4E**). Together these results indicate that the absence of collagen and therefore direct contact of bacteria with immune cells, causes a dysfunction in the neutrophil and macrophage response to infections resulting in bacterial persistence and an exaggerated inflammatory response.

**Figure 4.**
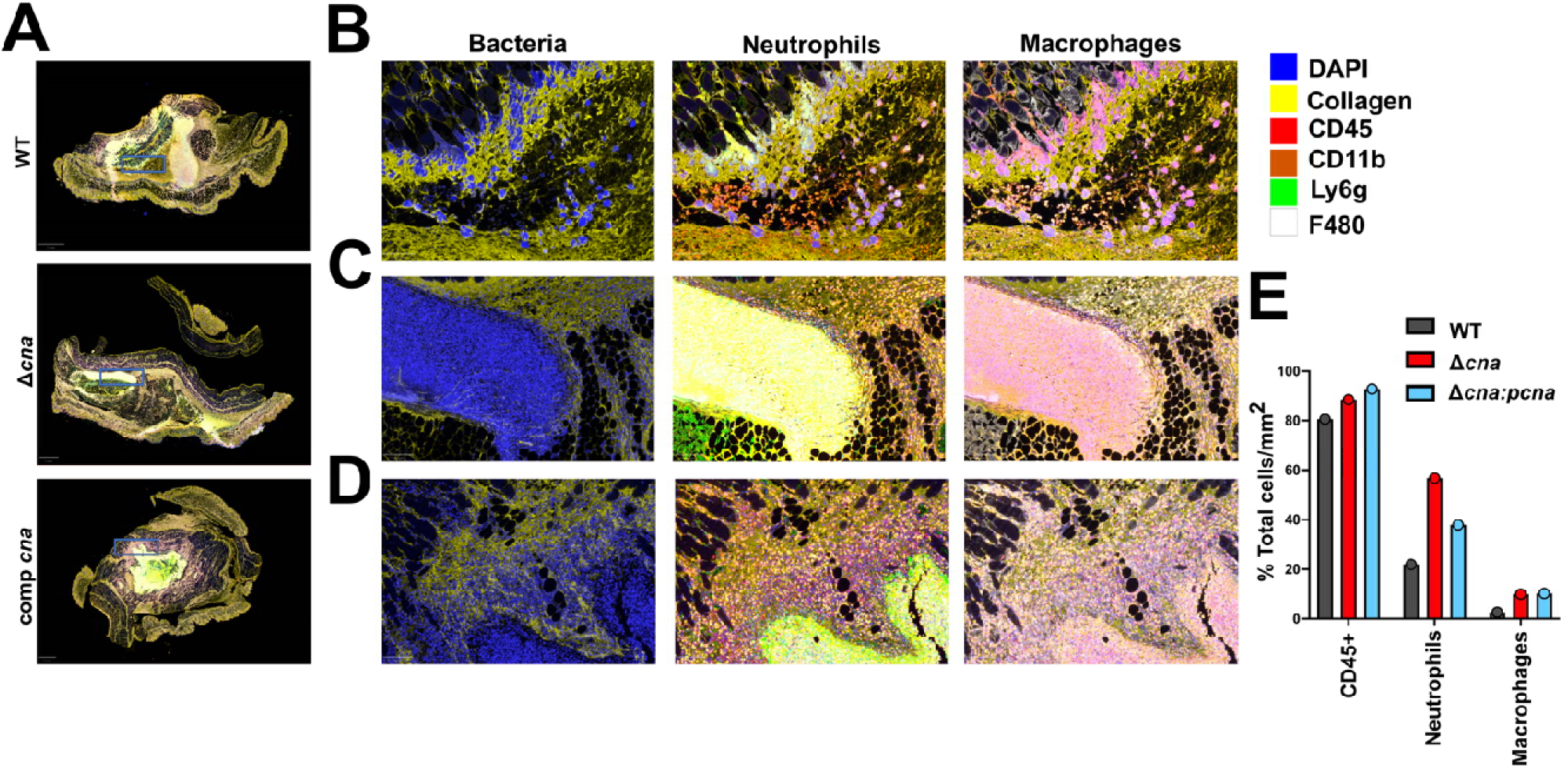
*S. aureus-* collagen aggregates trigger increased neutrophilic response in skin abscess infection. Multispectral quantitative pathology of skin abscess tissue excised from mice infected with WT, Δ*cna* or comp *cna* bacteria 3 days post inoculation and stained with antibodies targeting 6 host proteins as described in the legend. Images depict staining of entire tissue section as used for quantitative analysis (**A**). Images digitally zoomed in (X5.9) from sections shown in A. Regions of interest are depicted in A as blue boxes. Images demonstrate staining from tissues infected with WT (**B**) Δ*cna* (**C**) or comp *cna* (**D**) and stained with antibodies targeting 6 host proteins as shown in legend. Images are separated according to channels that demonstrate spatial distribution of bacteria with collage (DAPI, Collagen), neutrophils (CD45, CD11b) and macrophages (CD45, F480). Quantification of stainin demonstrated in A represented as total cells per millimeter of tissue sections infected with strains as described abov (**E**).

### Cna-C1q interaction dampens inflammation caused in response to *S. aureus in vivo*

C1q plays a central role in the opsonophagocytic response of neutrophils to bacteria^44^. Since C1q binds Cna in addition to collagen, we performed an experiment similar to (**Figure 3G-I)**, with bacteria that were opsonized with C1q depleted serum before incubation with human neutrophils. Under these conditions, we observed that neutrophils gained access to the collagen bound aggregates of WT and comp *cna* bacteria. This was associated with neutrophil lysis similar to the response generated to Δ*cna* bacteria in both C1q replete and depleted conditions. WT and comp *cna* were observed to elicit a response similar to Δ*cna* bacteria, in the absence of C1q (**Figure S4D-F**). To evaluate the contribution of the C1q-Cna interaction *in vivo*, we used the previously described abscess model of skin infection in C1q knockout mice (hereafter C1qKO) of the C57BL/6 WT background. Since our observations were made in the BALB/c mouse background, we performed a direct comparison of infection between C57BL/6 WT and C1qKO mice. Similar to previous results, we found that infection of C57BL/6 WT mice with Δ*cna* bacteria resulted in a ∼1.5 log increase in bacterial burdens (**Figure 5A**) and visually bigger abscess lesions that were significantly larger than both WT and comp *cna* infected animals when quantified (**Figure 5B**). We therefore measured the levels of inflammatory cytokines present in WT C57BL/6 abscesses and observed results similar to infections performed in female BALB/c mice (**Figure S5A**), with Δ*cna* infected abscesses containing significantly higher concentrations of key inflammatory markers when compared to both WT and comp *cna* groups (**Figure S5B**). To understand the role of the C1q-Cna interaction in this model we similarly infected C1qKO mice with WT, Δ*cna* or comp *cna* bacteria and observed an overall increase in the bacterial burden of WT and Δ*cna* infected animals retrieved at day 7, when compared with WT C57BL/6 infections (**Figure 5A**). Additionally, we observed a reduction in bacterial CFU loads (∼1 log) between WT and Δ*cna* infected mice. Similarly, the differences between WT and Δ*cna* lesions sizes were smaller in the C1qKO mouse background (**Figure 5B, C**). Lastly, the overall levels of inflammatory cytokines generated in the C1qKO infection background were higher and comparable between WT and Δ*cna* infected mice, once again demonstrating that C1q dampens inflammation in the presence of Cna (**Figure 5D, Figure S5C**).

**Figure 5.**
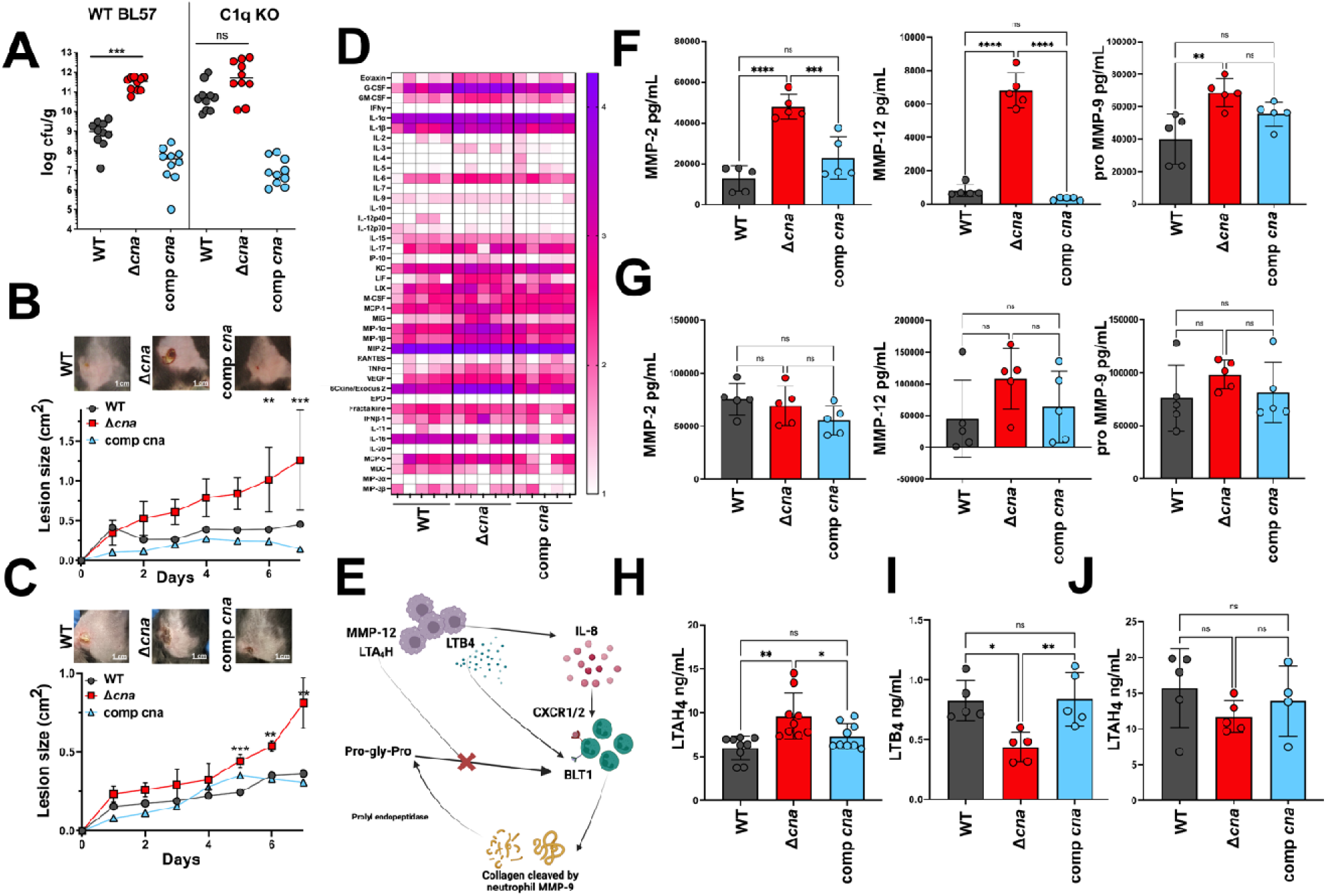
C1q-Cna binding controls matrix metalloprotease activity and inflammation in skin abscess. CFU per gram of homogenized tissue enumerated from either WT BL57 or C1qKO mice, 7 days post infection with WT, Δ*cna* or comp *cna* bacteria (**A**). Representative images of lesions formed at day 7 post infection of WT BL57 (**B**) or C1qKO (**C**) mice with strains described in A (top). Lesion sizes measured over the course of the infection for eac strain (n=10 per group, bottom). Comprehensive view for the concentrations of 44 inflammatory cytokines measured from abscess tissue 7 days after infection with respective bacteria, in the C1qKO mouse background. Concentrations are presented in logarithmic scale of picogram per mL homogenized tissue. Each column represents results from a single mouse (n=5 per group)(**D**). Graphical depiction of the known mechanisms by which matrix metalloproteases and 12 can instigate a cascade of neutrophil and macrophage mediated inflammation of skin infection sites (**E**). Concentrations of MMP-2, pro-9 and 12 measured from abscess tissue excised from WT BL57 (**F**) or C1qKO (**G**) mice infected with respective strains. Concentrations of leukotriene A-4 hydrolase (LTAH_4_) measured using a enzyme linked immunosorbent assay, from WT BL57 mice infected with bacterial strains as described in G (**H**). Assay similar to H measuring the concentrations of leukotriene B4 from homogenized tissue samples (**I**). Assay similar to H, quantifying the concentrations of LTAH_4_ and performed with tissue samples from C1qKO mice infected with bacterial strains as described in H (**J**). Results are representative of 2 independent analyses. Statistical analyses wer performed with a two-way ANOVA (B, C) or a one-way ANOVA (A, F-J) with Bonferroni posttest. Error was calculate based on SEM. *P<0.05, **P<0.01

Altogether our *in vitro* and *in vivo* results demonstrate that the absence of C1q increases severity of infection pathology and the Cna-C1q interaction results in WT *S. aureus* phenotypes that are similar to those observed in Δ*cna* infected animals.

### Tissue matrix metalloproteases contribute to macrophage dysfunction and excessive neutrophil influx

We observed a significantly higher level of the tissue inhibitor of matrix metalloprotease-1 (TIMP-1) measured from Δ*cna* infected mouse tissue, when compared to WT and comp *cna* groups at both day 3 (**Figure S3E**) and 7 (**Figure 2C, E**) post infection. This suggested that the host immune system may attempt to control the levels of inflammatory MMPs in the tissue bed during Δ*cna* infection. MMP-2 and 9 are gelatinases that also bind to collagen, while MMP-12 is a macrophage elastase^30,48^. MMP-9 is specifically released by neutrophils in response to the small molecule mediator, leukotriene <u>B</u>4 (LTB4), from macrophages^49^. MMP-9 degrades collagen to form the proinflammatory, neutrophil chemotactic peptide, Pro-Gly-Pro which functions by binding CXCR1/2, resulting in recruitment of additional neutrophils to the site of infection (**Figure 5E**). This would result in increased levels of the cytokine KC, similar to our observations at day 3 (**Figure S3E**) and 7 (**Figure 2E**).

Concentrations of MMP-2, 9 and 12 measured from abscess tissues were significantly higher in Δ*cna* infected tissues when compared to WT and comp *cna* groups (**Figure 5F**). Since both the C-terminal globular domain and N-terminal collagen tail of C1q are recognized by macrophages via the gC1qR and cC1qR/calreticulin receptors respectively, we asked whether this inflammatory cascade was disrupted in the absence of C1q. While C1qKO mice showed higher levels of all MMPs measured, when compared to WT mice, levels of MMPs in WT and comp *cna* infected groups were comparable with those measured from Δ*cna* infected mice (**Figure 5G**). To terminate this inflammatory cycle, proinflammatory, MMP-12-secreting macrophages release leukotriene A4 <u>h</u>ydrolase (LTAH4) which directly binds and inactivates Pro-Gly-Pro^33^. Δ*cna* infected mice contained significantly higher levels of LTAH4 compared with WT and comp *cna* groups (**Figure 5H**). Since LTAH4 is also required for the formation of LTB4, we reasoned that LTAH4 binding to increased levels of Pro-Gly-Pro would reciprocally limit the concentrations of released LTB4. Indeed, we observed a significant reduction in concentrations of this small molecule in Δ*cna* infected animals, compared with WT and comp *cna* infected mice (**Figure 5I**). Furthermore, these differences were found to be abrogated when LTAH_4_ was measured from C1q KO mouse infections (**Figure 5J**). Together these results indicate that the local cycle of inflammation caused by the direct interaction of macrophages and neutrophils with Cna negative *S. aureus,* is also tempered by the presence of C1q.

## Discussion

Studies characterizing the *cna* gene of *S. aureus* were performed over a decade ago^18,50^. Our knowledge of its importance to *S. aureus* pathogenesis, however, is extremely limited. Published reports were in three infection environments, namely eye, heart and bone, utilizing the less characterized strains of *S. aureus:* Phillips and CYL316^20,50,51^. These previous studies do not examine Cna in the context of skin colonization or SSTIs despite the essential role for collagen in the resolution of skin infection and the overwhelming presence of *S. aureus* in a majority of purulent SSTIs^52,53^. Here we provide a comprehensive analysis for the role of Cna, specifically during *S. aureus* skin infections. We demonstrate that the inability to directly bind collagen is associated with worsened outcomes **(Figure 1A-E)**, and that Cna-negative *S. aureus* is prevalent among major skin clones **(Figure 1 F)**.

Multiple comparative analyses that investigate the role of adhesins to inflammation caused during *S. aureus-* associated skin infection, do not identify Cna as an important virulence determinant^54^. In one study that included skin samples from patients with atopic dermatitis, psoriasis and normal skin colonized with *S. aureus*, absence of Cna expressed from *S. aureus* resulted in no observable decrease in bacterial binding to the stratum corneum^55^. Our studies demonstrate that Cna is sufficient and necessary to reduce the severity of skin infection caused by the predominant, clinically relevant, USA300 strain (**Figure 1G-J**). Further studies to specifically characterize the prevalence of isolates lacking the *cna* gene are therefore imperative to our knowledge of *S. aureus* skin infections. Of note, <u>P</u>anton <u>V</u>alentine leukotoxin (PVL) is epidemiologically linked to primary, purulent SSTIs ^56–58^. MW2 and USA300, the strains used in our study, both express PVL, indicating that the phenotypes observed here are likely not associated with the expression of this toxin.

Biochemical characterization of the Cna protein from strain Phillips, identifies the N-terminal A domain as the ligand binding region of the protein^16^. This domain can bind to complement protein C1q and prevent the opsonization of *S. aureus* ^22^. Authors in these studies used RBC lysis as a functional read out of complement activation. Here we additionally demonstrate opsonophagocytosis as a more relevant downstream effect **(Figure 3 A-C)**^59,60^. By using primary human neutrophils, we build on previous findings to show that when C1q is not sequestered by binding to Cna, this results in a likely uncontrolled mechanism of cell death with increased neutrophil lysis accompanied by bacterial survival (**Figure 6A, B**). Therefore, competition between collagen and C1q for binding to Cna correlates with the downstream neutrophil response, and the ability to fine tune ligand binding (collagen vs C1q) is necessary to protect *S. aureus* from immune clearance mechanisms^22^. Indeed, when Cna is ectopically expressed from a high copy expression vector, bacterial survival in C1qKO mice is comparable to WT and comp *cna S. aureus,* in WT C57BL/6 mice, indicating that increased binding to collagen may compensate for the absence of binding events that occur with the C1q N-terminal domain **(Figure 5A)**.

**Figure 6.**
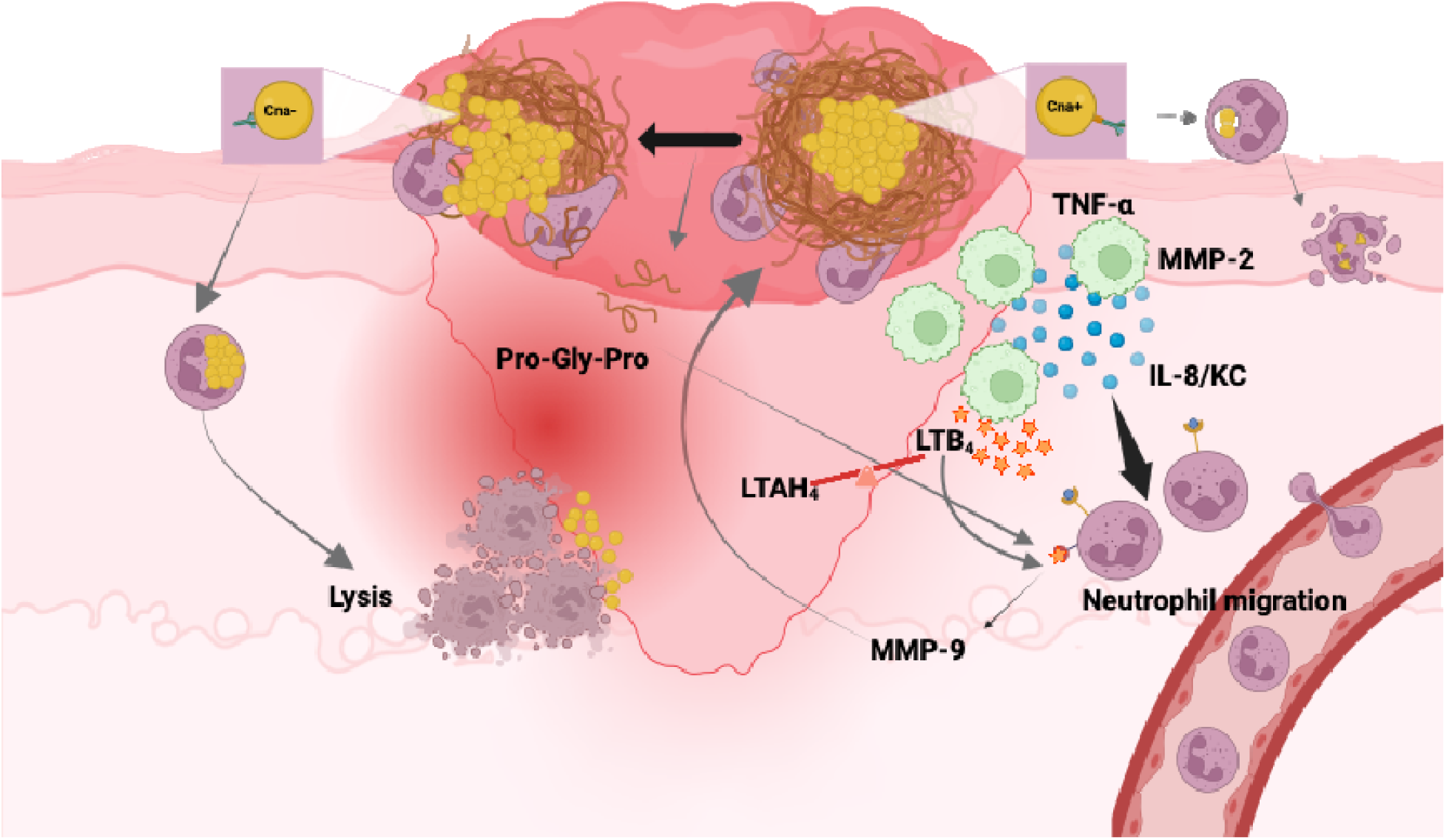
Summary of results. Expression of Cna allows *S. aureus* to bind to collagen and restrict bacterial contact with immune cells such as neutrophils. The N-terminal collagen like domain of C1q binds to Cna and reduces complement deposition and opsonophagocytosis by neutrophils. Bacteria that are taken up are eliminated by neutrophils (**A**). Bacteria lacking the ability to express Cna cannot directly bind to collagen. This allows direct contact between neutrophils and bacteria. C1q is not sequestered in the absence of Cna, causing increased bacterial uptake by neutrophils. This leads to neutrophil lysis and inflammation (**B**). Macrophages in the infection bed will releas proinflammatory mediators including the matrix metalloprotease MMP-2, MMP-12 and neutrophil chemokine, IL-8. Macrophages also express leukotriene A4 hydrolase (LTAH_4_) in response to infection, which activates the solubl effector, leukotriene B 4 (LTB_4_). Neutrophils release MMP-9 which breaks down collagen to inflammatory Pro-Gly-Pr which assists in further neutrophil recruitment (**C**). Image created using Biorender.com

It is likely that while Cna-C1q binding affects opsonization and uptake of bacteria as shown here, toxins released by *S. aureus* are responsible for bacterial escape and neutrophil lysis. Leukotoxins are widely characterized as causing immune cell lysis, with LukAB being particularly significant at allowing intracellular bacteria to lyse neutrophils from within ^61–63^. This activity occurs in concert with alpha hemolysin (Hla), a secreted, pore forming toxin^64,65^.Further studies that expand on the effect of Cna and its interactions with host ligands, with emphasis on the neutrophil response to *S. aureus,* are required in order to further evaluate potential roles for these toxins.

Our results provide additional evidence for the vital role that macrophages play during resolution of skin infections (**Figure 6C**)^66^. In this work, the absence of viable macrophages was accompanied by increased numbers of dead neutrophils and prolonged inflammation in mice infected with Δ*cna* bacteria, providing further validation of the communication between neutrophils and macrophages that is essential for resolution of skin infection. Hla has documented roles in macrophage lysis, leading to reduced neutrophil infiltration and bacterial clearance during skin infection ^67^. Increased bacterial survival in the absence of Cna may result in higher local concentrations of Hla and therefore more lysed macrophages. Whether Hla plays an additional role in the lack of live macrophages observed in our studies, remains to be assessed and would provide a deeper understanding of the triggers that cause macrophage activation, small molecule release and therefore neutrophil infiltration. Similarly, the increased concentrations of matrix metalloproteases may be a direct effect of the presence of a larger number of lysed neutrophils. One report indicates that matrix metalloproteases 1, 2, 3 and 9 can cleave the collagen-like domain of C1q and result in neutrophil reactive oxygen burst. This may contribute to the killing of bacteria that are exposed to neutrophils **(Figure 4 G-I)**^68^. Lastly, while we see a rise in observable, dead neutrophils when mice are infected with Δ*cna* bacteria, we do not observe these differences in macrophage populations at the timepoints examined (**Figure 2J**). Macrophages can suffer numerous fates following resolution of inflammation including conversion to endothelial cells or fibroblasts^69^. Recently, fibroblasts have been demonstrated to be an important source of collagen that is utilized as nutrition by *S. aureus* during pulmonary infections. Of note, these studies were performed in strains that do not express Cna^70^. The fate and function of macrophages and monocytes following exposure to *S. aureus* in the context of Cna, remains to be resolved ^71,72^.

Results presented here translate *in vitro* experiments for the first time to demonstrate a direct association between the Cna-C1q interaction and control of inflammation *in vivo* (**Figure 5A-D**). It is important to note however, that C1q is one of many host proteins that contain a collagen-like motif, any of which could be present in the abscess microenvironment and contribute to sequestration of available ligand binding domains on Cna^73,74^. Additionally, host proteins that bind the N-terminus of C1q, such as MBL-associated serine proteases, could compete with Cna for binding^75^. Similarly, most studies focus on the ability of Cna to bind type 1 collagen. The ratio of type 1/3 collagen is crucial for wound healing and most matrix metalloproteases bind multiple types of collagens. This may influence the outcome of infection in the context of Cna, and remains to be studied^48,76^.

USA300 (Cna-) and MW2 (Cna+) are both successful, clinically isolated strains of *S. aureus* that differ in their ability to express Cna^7,13,77^. This, together with our findings indicates that rather than determining bacterial survival, the expression of Cna may dictate the nature and length of infection caused by *S. aureus,* as well as the degree of inflammation achieved as a consequence.

Our current work reveals that rather than assisting in infection exacerbation and dissemination of *S. aureus* such as would be expected of a canonical virulence factor, Cna is involved in concealing bacteria from the immune system and quiescently establishing an infection bolus, likely until favorable conditions for growth become unavailable and dissemination is required for survival. These findings also indicate that the loss of *cna* may have provided an evolutionary advantage to make *S. aureus* prolific at dissemination^78,79^. Conversely, expression of Cna by a subset of strains may allow them to persist for long periods in the community, either as colonizers or as chronic, biofilm associated infections ^20,50,51^.

*S. aureus* cells expressing Cna accumulate collagen which may serve as a ’self’ signal that allows immune evasion. Bacteria afford the time required to proliferate and form a collagen shield by binding to collagen like domains of the major innate, bacterial recognition protein, C1q. This function potentially extends to additional pathogen recognition molecules, many of which express similar domains^73,74^. Additionally, bacteria that are recognized and engulfed by immune cells presumably utilize one or more bacterial toxins to kill these cells, allowing *S. aureus* to survive^47,80,81^. We demonstrate that the loss of *cna* promotes the expansion of *S. aureus* infection. It is well established that when *S. aureus* is present in sufficient numbers, it is not easily eliminated by the immune system^47,80,82^. Our results with skin infections confirm this and show that this results in massive immune cell death, likely caused due to bacterial virulence properties that are similar to Cna expressing cells. The zone of necrotic cells, largely neutrophils, presumably shields bacteria from the entry and function of additional immune cells into the infection bolus. Altogether, this work establishes a major role for Cna in *S. aureus* skin infections and demonstrates its significant immune evasion properties.

## Resource availability

### Lead Contact

Further inquiries and information on reagents and resources should be directed to (and will be fulfilled by) the lead contact, Alexander R. Horswill.

### Materials availability

Reagents and materials used or generated in this study can be made available upon request from the lead contact.

### Data availability

Data reported in this manuscript will be made available by the lead contact upon request.

## Acknowledgments

The authors would like to thank members of the Horswill and Doran groups for their critical evaluation of the data in this manuscript. We also acknowledge the Human Immune Monitoring Shared Resource (RRID:SCR_021985) within the University of Colorado Human Immunology and Immunotherapy Initiative and the University of Colorado Cancer Center (P30CA046934) for their expert assistance in analysis of multispectral quantitative pathology. We thank Laura Hoaglin HT(ASCP) with the Gates Histology Services Core Lab (University of Colorado Anschutz Medical Campus) for microtomy and hematoxylin-eosin staining. B.S is funded by the <u>N</u>ational Institutes of <u>H</u>ealth (NIH) awards R01AI137336 and R01AI140754. K.S.D is funded by NIH awards R01NS116716 and R01AI153332. A.R.H is funded by NIH award AI083211 and the Department of Veteran’s Affairs award BX002711.

## Author Contributions

Conceptualization, M.B., J.M.K., A.R.H.; Methodology, M.B., B. L.S., J.M.K., M.P., G.P., A.R.H; Investigation, M.B., B.L.S., J.M.K; Writing-Original Draft, M.B., Writing-Review and Editing, M.B., B.L.S, J.M.K., B.S., K.S.D., A.R.H.; Funding Acquisition, B.S., K.S.D and A.R.H.; Supervision, A.P., B.S., K.S.D., A.R.H.

## Supplemental Information

Document S1. Figures S1-S5

## Ethics Statement

Experiments with animals were reviewed and approved by the institutional animal care and use committee at the University of Colorado Anschutz Medical Campus (IACUC #00486) Primary human neutrophils were isolated from healthy human donors after obtaining informed, written consent from each donor. This was done according to the protocol approved by the University of Colorado Anschutz Medical Campus institutional review board (IRB #17-1926).

## Materials and Methods

### Bacterial strains and growth conditions

Unless otherwise indicated, all strains of *Staphylococcus aureus* were grown in tryptic <u>s</u>oy <u>b</u>roth (TSB) at 37°C with shaking. Overnight bacterial suspensions were sub-cultured and grown to exponential phase for all *in vitro* assays. This corresponded to an optical density of 0.42 (O.D. 600). Antibiotics were added during growth where indicated. *E. coli* ER2566 was grown in Luria Bertani Broth using methods described below for protein purification.

### Generation of bacterial mutants and complementation

Chromosomal deletion of the *cna* gene were performed using previously established methods^83^. Briefly, the temperature sensitive pJB38 plasmid was used to introduce DNA fragments (∼1kb) flanking the target region of interest. Flanking DNA was amplified (Phusion high fidelity polymerase, NE Biolabs) using gene specific primers, products were digested with restriction enzymes and purified (Qiagen PCR purification). Following triple ligation into pJB38, the plasmid was electroporated into *E. coli* DC10B and selected for on Luria Bertani agar plates containing 100 µg/mL ampicillin. Following confirmation from single colonies, plasmid was purified, PCR used for confirmation with construction and sequencing primers performed and plasmid was electroporated into *S. aureus*. Positive clones were selected on tryptic <u>s</u>oy <u>a</u>gar (TSA) containing 10 µg/mL chloramphenicol and homologous recombination performed at 42°C for 24 hours. Following overnight incubation in TSA-Cam and a series of subcultures in TSB at 30°C, counterselection was performed on 200 ng/mL anhydrotetracycline (30°C/overnight). Loss of plasmid was indicated by growth on TSA but not TSA-Cam and presence of desired mutation was verified using PCR with chromosomal primers that were outside the region of mutation. The *cna* complementing plasmid was generated by amplifying *cna* with its promoter region using the Q5 polymerase from WT MW2 genomic DNA using the primers:

F-ctcggtaccttaggaggatgattatttatgaacaagaacgtgttgaa

R-acagctatgacatgattacgaattcttatgagttaaatctttttcttaaaattaaatac

KpnI and EcoRI were used to digest this fragment which was subsequently ligated into pCM28 digested with the same enzymes. Plasmid was confirmed and maintained with 10 µg/mL chloramphenicol.

### Murine skin abscess model of infection

All infections were performed on 7-week-old mice by inoculating 1X10^8^ CFU/mL bacteria intradermally as previously described^34,35^. Briefly overnight bacterial cultures were sub-cultured and grown to an appropriate OD and resuspended in saline. Mouse stomachs were shaved and treated with Nair, hair removal cream, one day prior to inoculation of bacteria. Abscess formation was monitored with imaging over the span of 7 days. Mouse weight loss was measured daily. On day 7, mice were sacrificed, and abscesses excised using a 6mm punch biopsy. Abscess tissue was resuspended in 500uL phosphate buffered saline and homogenized with physical disruption using (0.1mm beads, Biospec Mini-Beadbeater). Colony forming units were calculated from homogenate and plotted per gram of tissue.

### Detection of *cna* from clinical isolates

We assessed the presence of *cna* by analyzing the genome sequences of isolates representing the major clonal types of methicillin-susceptible and -resistant *S. aureu*s isolates from two previously published studies of primary purulent SSTI and nasal colonization^36,37^. Briefly, libraries were prepared and sequenced at the NYU Langone Genome Technology Center using an Illumina NovaSeq to produce paired-end 150 bp reads. Reads were filtered and trimmed with fastp v0.20.1 using default settings^84^. Confindr v0.7.4 identified within-species contamination, and isolates with >10% contamination were excluded^85^. Filtered reads were then assembled with Unicycler v0.4.8 in conservative mode^86^. Taxonomic classification of assemblies was performed using GTDBTK v1.5.1; non–*S. aureus* isolates were excluded^87^. *S. aureus* sequence types and clonal complexes were determined with MLST (https://github.com/tseemann/mlst). We used BLAST v2.12.0+ to search for the *cnaB* gene with the KEGG sequence (ID: MW2612), considering a genome *cnaB* positive if a BLAST hit showed an E-value < 10 ² ^88,89^.

### Mouse histology

Abscess tissue was fixed in 10% formalin and submitted to the Gates Histology Services Core Lab (University of Colorado Anschutz Medical Campus) for microtomy and hematoxylin-eosin staining.

### Cytokine measurements

Cytokine concentrations were measured from mouse tissue homogenate that was treated with protease inhibitor (Sigma) and stored in PBS at -80°C before being analyzed by Eve Technologies. Briefly the multiplexing analysis was performed using the Luminex™ 200 system (Luminex, Austin, TX, USA) by Eve Technologies Corp. (Calgary, Alberta). Forty-five markers were simultaneously measured in the samples using Eve Technologies’ Mouse Cytokine 45-Plex Discovery Assay®. Assay sensitivities of these markers range from 0.3 – 30.6 pg/mL for the 45-plex. Individual analyte sensitivity values are available in the MilliporeSigma MILLIPLEX® MAP protocol.

### Murine skin abscess flow cytometry

Murine model of skin abscess infection with *S. aureus* was performed as described above. Skin abscesses were harvested using a 6mm biopsy punch following perfusion of the animal with PBS and heparin. Skin punches were minced and incubated for 2 hours at 37 C with shaking in RPMI 1640 with Miltenyi Multi Tissue Dissociation Kit 1 enzymes (volumes according to the manufacturer). Following enzymatic digestion, samples were mechanically digested using the Miltenyi Gentle Macs dissociator (program “Multi H”). Single cell suspensions were prepared from digested abscess samples by filtering samples through a 70 mm cell strainer and pelleting cells by centrifugation (300 x g, 5 minutes). Remaining red blood cells were removed by resuspending the cell pellet in red blood cell lysis buffer (150 mM NH_4_Cl, 10 mM KHCO_3_, 0.1 mM Na_2_EDTA; pH 7.2) for 2 minutes at room temperature and washing with RPMI 1640 (Gibco). Cells were then pelleted by centrifugation and resuspended in MACS buffer (Phosphate buffered saline with 0.5% BSA and 2 mm EDTA, pH 7.2). Single cell suspensions were first stained with eBioscience Fixable Viability Dye eFluor 506 (Catalog # 65-0866-18) in PBS for 30 minutes at room temperature. Cells were stained with the anti-mouse surface antibodies in MACS buffer for 30 minutes at room temperature (See Supplemental Figure 2B for details on antibodies). After surface antibody staining, the cells were fixed (30 minutes at room temperature) using the FoxP3 fixation/permeabilization kit (Thermo Fisher Scientific, Catalog # 00-5523-00). Stained cells were analyzed on a BD LSRFortessa (BD Biosciences) using the BD FacsDiva software (v9) or Cytoflex. Data were analyzed with BD FlowJo software v 10.10.0. Gating strategy was adapted from a previous study and can be found in Supplemental Figure 2A^42^.

### C4b and C3b binding assays

To measure the differences in C3 binding between various strains, bacteria were grown overnight in TSB and brought up to an O.D of 0.42 were washed 3 times with PBS and re-suspended in 10% pooled human serum for 30 minutes. After repeating the washing step, each sample was stained with rabbit monoclonal Alexa Fluor 647-labelled anti-C3 antibody (Abcam ab196639) or FITC labeled anti-C4b antibody (Thermo Fisher, PA1-28407) for 30 or 20 minutes respectively, at room temperature. Samples were then washed 3 times with PBS and resuspended in PBS for analysis by flow cytometry, using the BD LSR Fortessa (BD Biosciences) and BD FacsDiva software (v9).

### Neutrophil isolation

Isolation of primary human neutrophils was performed using whole blood collected from healthy, consenting human volunteers. Methods used are as previously described^46^. Briefly, whole blood components were separated using a Ficoll Hypaqe based density gradient. Following removal of peripheral blood monocytes and lymphocytes, red blood cells were lysed with water and remaining neutrophils were resuspended in 0.9% sodium chloride. Following re-suspension in phosphate buffered saline neutrophils were enumerated using a hemocytometer. Neutrophils were used at a concentration of 4X10^6^cells/mL as previously published^46^.

### Neutrophil opsonophagocytosis assays

*S. aureus* was incubated with primary, blood-derived human neutrophils isolated as previously described^46^. To do this, overnight cultures of bacteria grown in tryptic soy broth were diluted 1:100 into fresh medium and grown to an O.D of 0.42, corresponding to ∼1X10^8^ CFU/mL. Cultures were then washed 3 times with phosphate buffered saline (PBS) and re-suspended in 10% pooled human serum (Complement Tech) for 30 minutes, in order to opsonize bacteria. Cultures were washed 3 times with PBS and resuspended in 4X10^6^ neutrophils/mL, to bring the multiplicity of infection to 1 (neutrophil): 25 (Bacteria). Samples were then incubated at 37 degrees for 10 (uptake) or 30 (survival) minutes and treated with 10ug/mL lysostaphin for 5 minutes to eliminate extracellular bacteria, as previously described^45^. Intracellular bacterial survival was assessed by plating serial dilutions on tryptic soy agar. Percent survival was calculated by comparing survival for each strain at 10 and 30 minutes, to its bacterial inoculum present at time 0.

### Neutrophil confocal microscopy

Bacterial strains were opsonized as described above either with 20 μg/ml type I rat tail collagen (Corning®, Cat No: 354236), or without collagen as a control. Staining was performed using previously published methods^47^. Briefly, neutrophils were isolated as described above (4X10^6^/mL) and stained with Cell Tracker Blue CMAC (Thermo Fisher, Cat No: C2110). Opsonized bacteria labelled with Syto-9 (Invitrogen, Cat No: S34854) were washed and incubated with neutrophils (1:25) for 20 minutes, similarly to method described above. Samples were washed (X3 PBS) and centrifuged (800g) for 15 minutes, pellets re-suspended in minimal volume and mounted onto glass slides with Pro Long Gold Antifade (Invitrogen, Cat No: P36930) with coverslips. Images were collected using the Olympus FV1000 confocal laser scanning microscope (Advanced Light Microscopy Core, University of Colorado Anschutz Medical Campus).

### Matrix metalloproteinase measurements

MMP concentrations were measured from mouse tissue homogenate that was treated with protease inhibitor and stored in PBS at -80°C before being analyzed by Eve Technologies. Briefly, the multiplexing analysis was performed using the Luminex™ 200 system (Luminex, Austin, TX, USA) by Eve Technologies Corp. (Calgary, Alberta). Five markers were simultaneously measured in the samples using Eve Technologies’ Mouse MMP 5-Plex Discovery Assay® (MilliporeSigma, Burlington, Massachusetts, USA) according to the manufacturer’s protocol. The 5-plex consisted of MMP-2, MMP-3, MMP-8, proMMP-9 and MMP-12. Assay sensitivities of these markers range from 1.6 – 8.4 pg/mL for the 5-plex. Individual analyte sensitivity values are available in the Millipore Sigma MILLIPLEX® MAP protocol

### Measurement of LTAH_4_ and LTB_4_ from abscess tissue

Enzyme linked immunosorbent assays were performed to measure the concentration of LTAH_4_ (Biomatik, Cat No: EKN46742-96T) and LTB_4_ (Avantar, Cat No: 76576-968) according to the manufacturer’s protocol. Quantification was done using abscess tissue that was biopsied on the same day (day 7 post infection), homogenized as described above and re-suspended in PBS to be used immediately.

### Immunofluorescence staining

Performed on paraffin embedded tissue sections excised from 7-week-old, female BALB/c mice which were euthanized 3 days post inoculation with WT, Δ*cna* or comp *cna* bacteria, or saline as a negative control. Analysis done by the Human Immune Monitoring Shared Resource, University of Colorado Cancer Center

