## Supplemental Figures for "Collagen binding adhesin restricts *Staphylococcus aureus* skin infection"

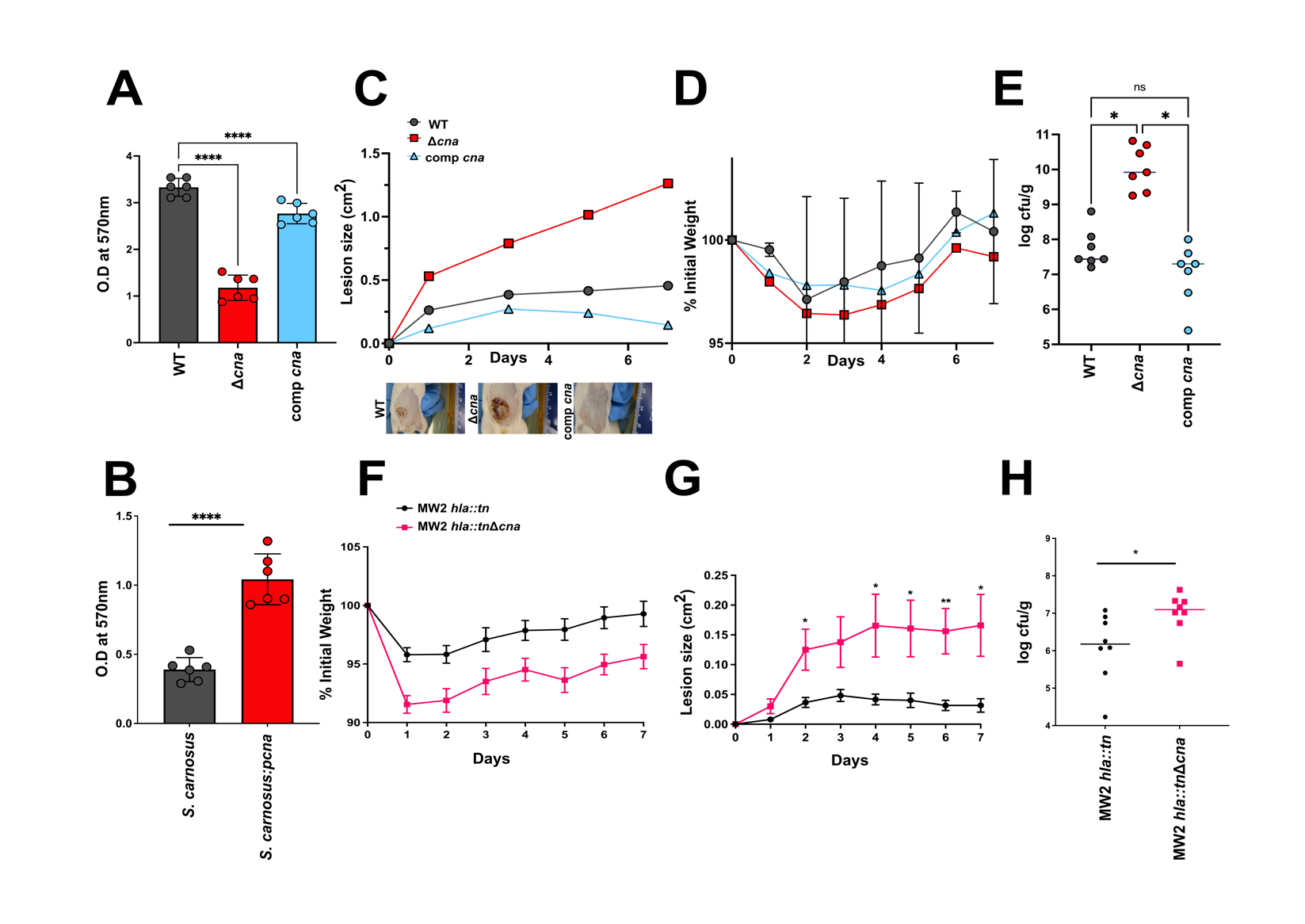
**Figure S1. Collagen binding adhesin is sufficient to reduce infection severity in a sex independent manner, related to Figure 1.** Crystal violet-based adhesion assay to compare the collagen binding capacity of WT, isogenic *Δcna* or comp *cna* bacteria on a collagen-coated surface (**A**). Adhesion assay similar to A, performed with *Staphylococcus carnosus* (Cna-) or *S. carnosus:pcna* (Cna+) (**B**). Lesions sizes measured from male Balb/c mice (n=7 per group) infected with strains mentioned in A, 7 days post inoculation. Measurements were made using Image J. Images are representative of lesions formed by each strain at day 7 (**C**). Weight loss measured from mice infected as described above , over 7 days and calculated as a percent of weight measured at day 0 (**D**). CFU per gram of homogenized tissue collected at day 7 post infection with strains described in C (**E**). Weight loss measured similar to C, in female Balb/c mice (n=8) infected with MW2 *hla::tn* lacking alpha hemolysin expression, or the isogenic MW2 *hla::tnΔcna* strain, plotted as a percent of the weight recorded at day 0 (**F**). Lesion sizes measured from mice infected as described in F, calculated using Image J (**G**). CFU per gram of tissue enumerated at day 7 post infection in mice infected with strains as described in F (**H**). Results are representative of 2 independent analyses. Statistical analyses were performed with a one-way ANOVA (A, E) a two tailed Students t-test (B, H) or a two-way ANOVA with Bonferroni post test. Error was calculated based on SEM. *P<0.05, **P<0.01, ****P<0.0001

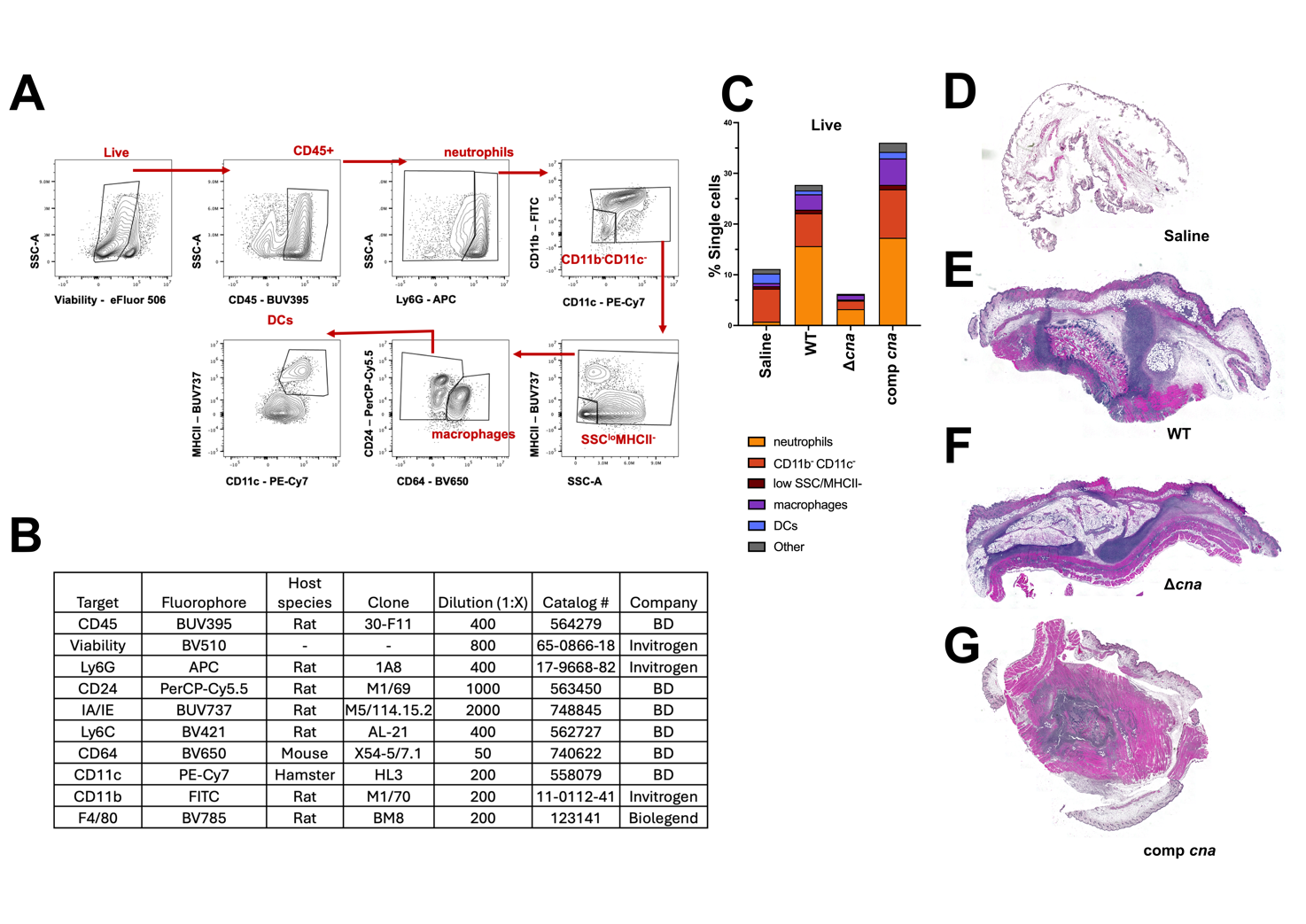
 **Figure S2. Assessing the inflammatory response to *S. aureus* collagen binding adhesin, related to Figure 2.** Demonstration of gating strategy utilized to perform flow cytometry on excised abscess tissue. Sample shown is tissue infected with WT MW2 bacteria and collected at day 3 post inoculation **(A).** Antibody panel used to measure immune cell types present in infection as shown in A, **(B).** Summary of results from flow cytometry performed on excise abscess tissue from mice that were infected with WT, *Δcna* comp *cna* bacteria for 7 days (n=7 per group). Results show percentages of total single, live cells subdivided according to gates set as in A **(C)**. Hematoxylin Eosin staining on tissues collected at day 3 post intradermal inoculation of female Balb/c mice with saline **(D)** WT MW2 **(E)**, *Δcna* **(F)** or comp *cna* **(G)** bacteria. Results are representative of n=10 mice per group.

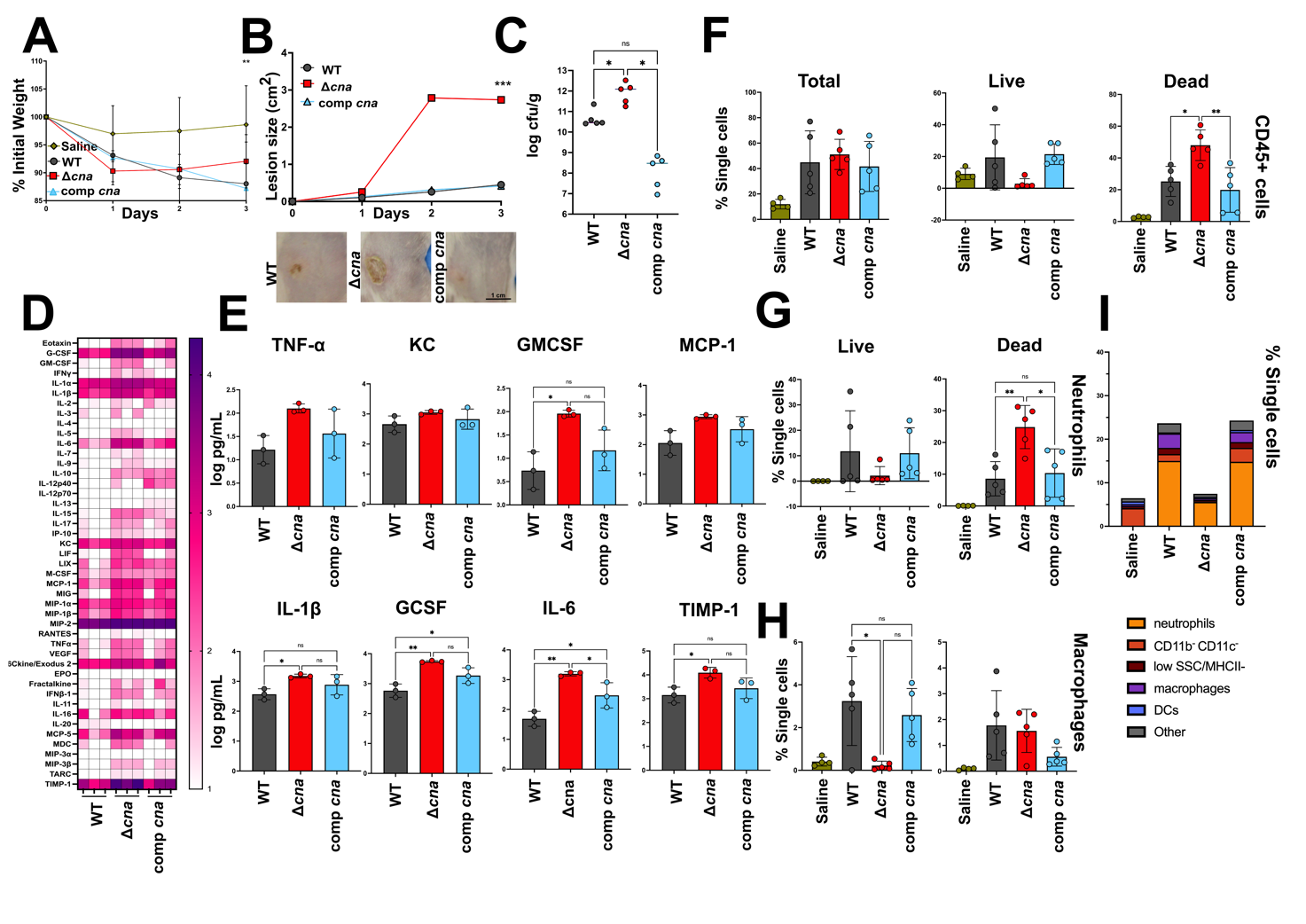
**Figure S3. Inflammation caused in the absence of Cna begins early in infection, related to Figure 2.** Weight loss measured from mice infected as described above, 3 days post inoculation with WT, *Δcna* or comp *cna* (n=5 per group). Percentage weight loss is measured in comparison to values at day 0 **(A)**. Lesion size measured at day 3 post infection, performed as described for A. Measurements were made using Image J **(B)**. CFU per gram enumerated from abscess tissue biopsied from mice infected as described in A (n=5 per group) **(C)**. Summarized view for the concentrations of 44 inflammatory cytokines measured from abscess tissue 3 days after infection with respective bacteria. Concentrations are presented in logarithmic scale of picogram per mL homogenized tissue. Each column represents results from a single mouse (n=3 per group) **(D).** Individual graphs demonstrating the concentrations of 7 markers of inflammation as well as the tissue inhibitor of metalloproteases, as measured using the assay described in D **(E)**. Flow cytometry quantification of CD45+ cells from abscess tissue collected 3 days post infection with strains as described in A or saline as a control **(F)**. Quantification of numbers of live and dead neutrophils **(G)** and macrophages **(H)** observed in tissue samples collected similar to A. Populations were identified using gating strategy described in Supplementary Figure 2A. Graphical summarization of all measured CD45^+^ populations (see Supplementary Figure 2A) from abscess tissue collected at day 3 post infection with respective strains or saline as a negative control **(I).** Statistical analyses were performed with a two-way ANOVA (A, B) or a one-way ANOVA with Bonferroni posttest. Error was calculated based on SEM. *P<0.05, **P<0.01

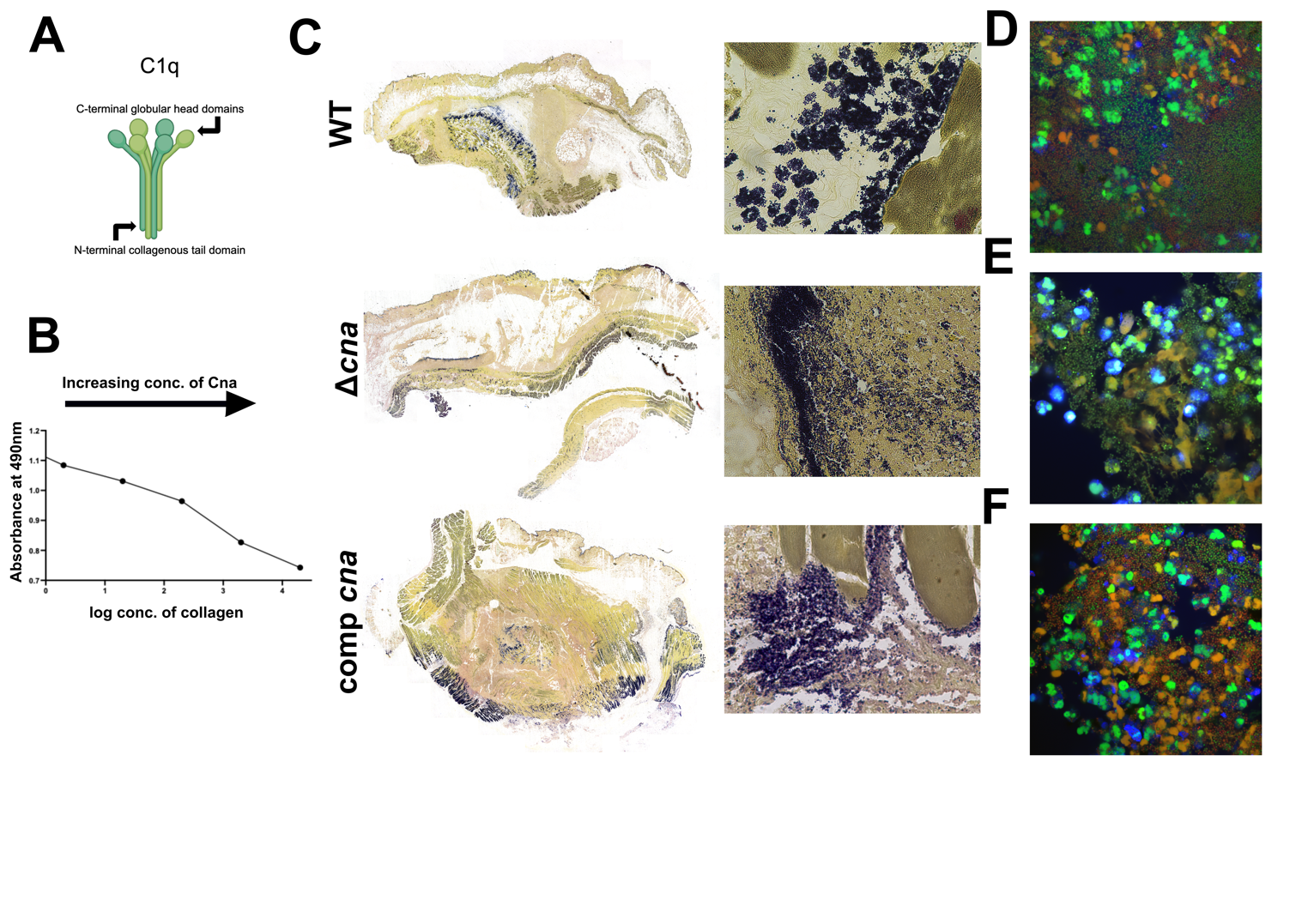
**Figure S4. Cna binds to collagen motifs to localize infection and impede neutrophil phagocytic response, related to Figures 3, 4.** Graphical representation of the structure of serum C1q protein (**A**). ELISA performed to measure the binding of a range of concentrations of recombinant Cna to type 1 collagen following incubation with C1q (**B**). Modified Gram staining performed on tissue sections excised from mice at day 3 post intradermal infection with WT, *Δcna* or comp *cna* bacteria (left) with region of interest from figure 4 shown with a digital zoom of X5.9 (right) (**C**). Confocal microscopy performed on respective bacterial strains (Green=Syto-9/live) opsonized with C1q-depleted pooled human serum, in the presence of type 1 collagen and incubated with Cell Tracker Blue-labelled primary human neutrophils for 20 minutes, following which samples were stained with ethidium homodimer-1 to visualize dead/dying cells (red) (**D-F**). Images were captured at X40 (A-C) and X1000 (D-F) magnification.

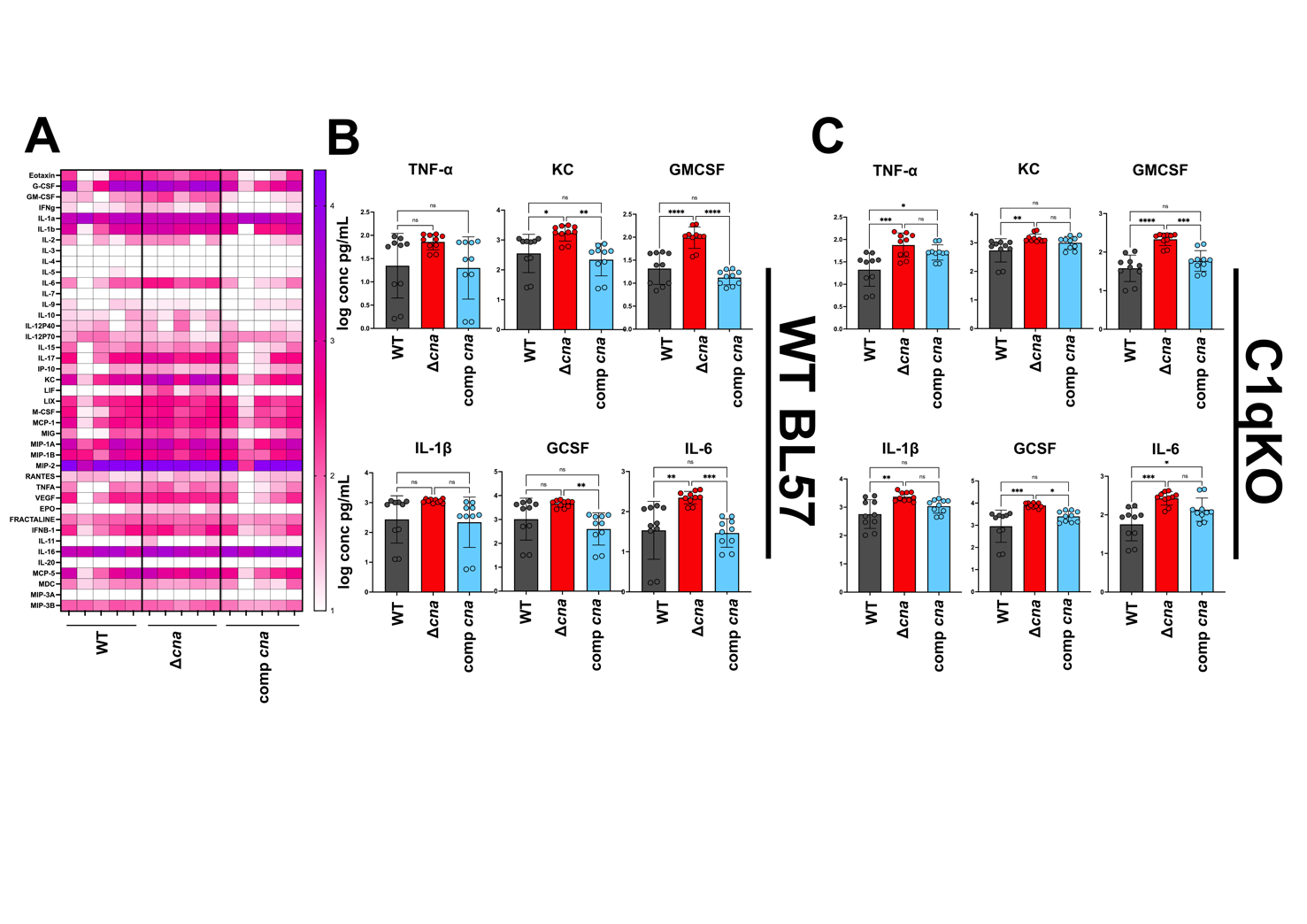
**Figure S5. Cna- C1q interaction dampens release of inflammatory cytokines in response to *S. aureus* intradermal infection, related to Figure 5**. Summarized view for the concentrations of 45 inflammatory cytokines measured from abscess tissue 7 days after infection with respective bacteria, in the WT BL57 mouse background. Concentrations are presented in logarithmic scale of picogram per mL homogenized tissue. Each column represents results from a single mouse (n=5 per group) (**A**). Individual graphs for 6 of the cytokines measured in A, measured from mice infected with strains as described for A (**B**). Individual graphs similar to B taken from cytokine multiplex array done in Figure 5, for infections done in the C1qKO mouse background (**C**). Statistical analyses were performed with a one-way ANOVA and Bonferroni post-test. Error was calculated based on SEM. *P<0.05, **P<0.01, ****P<0.0001
